# A highly sensitive strand-specific multiplex RT-qPCR assay for quantitation of Zika virus replication

**DOI:** 10.1101/2022.04.22.489195

**Authors:** Trisha R. Barnard, Alex B. Wang, Selena M. Sagan

## Abstract

Reverse-transcription quantitative polymerase chain reaction (RT-qPCR) is widely used to quantify viral RNA genomes for diagnostics and research, yet conventional RT-qPCR protocols are unable to accurately distinguish between the different viral RNA species that exist during infection. Here we show that false-priming and self-priming occur during reverse transcription with several published Zika virus (ZIKV) primer sets. We developed a RT-qPCR assay using tagged primers and thermostable reverse transcriptase, which greatly reduced the occurrence of nonspecific cDNA products. Furthermore, we optimized the assay for use in multiplex qPCR which allows for simultaneous quantitative detection of positive-strand and negative-strand ZIKV RNA along with an internal control from both human and mosquito cells. Importantly, this assay is sensitive enough to study early stages of virus infection *in vitro*. Strikingly, using this assay, we detected ZIKV negative-strand RNA as early as 3 h post-infection in mammalian cell culture, at a time point prior to the onset of positive-strand RNA synthesis. Overall, the strand-specific RT-qPCR assay developed herein is a valuable tool to quantify ZIKV RNA and to study viral replication dynamics during infection. The application of these findings has the potential to increase accuracy of RNA detection methods for a variety of viral pathogens.

**Highlights:**

- Self-primed cDNA is amplified by widely-used ZIKV qPCR primer sets
- Use of tagged primers and thermostable RT increases strand-specificity for RT-qPCR
- Multiplexed qPCR allows for simultaneous quantitation of (+) and (-) strand viral RNAs, and an internal control
- Strand-specific RT-qPCR can detect fewer than one copy of viral RNA per cell in human and mosquito cells

## 1. Introduction

Zika virus (ZIKV) is a mosquito-borne positive-sense RNA virus and member of the *flavivirus* genus in the *Flaviviridae* family (1). ZIKV infections are causally linked with congenital neurological complications, and ZIKV has caused a series of outbreaks of increasing severity in the past 15 years (2). Like all positive-sense RNA viruses, ZIKV replication proceeds through a negative-strand replication intermediate. New positive-strands are synthesized from the negative-strand template, and positive-strand synthesis generally outnumbers negative-strand synthesis by ∼10-100 fold (3-6). Negative-strand RNA detection is therefore the gold standard for detection of ZIKV replication.

Despite being a widely used method for quantification of viral RNA, standard reverse transcription quantitative polymerase chain reaction (RT-qPCR) protocols are unable to distinguish between the multiple species of viral RNA present in a sample. As a result, standard RT-qPCR protocols are unable to determine the absolute quantity of viral genomes due to the presence of both positive- and negative-strand viral RNA (7-10). False priming of the incorrect strand, self-priming by secondary structures in the viral RNA template, or random priming by contaminating nucleic acids have been proposed to contribute to the lack of strand-specificity in standard RT-qPCR assays (11-16). Strategies to improve specificity of RT-qPCR have been developed for multiple RNA viruses, and typically involve one or more of the following: 1) use of tagged RT primers containing a unique non-viral “tag” sequence at the 5’ end of a viral-specific sequence, and the use of tag-specific primers in qPCR; 2) high-temperature RT to minimize RNA template secondary structure, or the use of a reverse transcriptase with increased specificity; or 3) purification of complementary DNA (cDNA) products to avoid excess primer carry-over into the qPCR reaction (7, 8, 10, 17). Tagged primers have been successfully used for the strand-specific detection of some members of the *Flaviviridae* family (6, 7). However, previously-published assays for ZIKV negative-strand detection either do not use tagged primers (18-23) or have not demonstrated adequate strand-specificity (24-28). Additionally, although the use of DNA hydrolysis probes has been shown to improve the dynamic range of strand-specific qPCR, many previously-published strand-specific assays use intercalating dye chemistry (e.g. SYBR) for real-time detection in qPCR. Intercalating dyes detect all double-stranded nucleic acid non-specifically, and therefore multiple targets cannot be distinguished in a single PCR reaction (29-31). In contrast, detection of PCR amplification using hydrolysis probes can enable the detection of multiple targets from the same sample and would therefore allow for simultaneous detection of positive- and negative-strand viral RNAs, with normalization to an internal control.

Herein, we show that conventional RT-qPCR of ZIKV RNA generates cDNA from self- and false-priming. We show that both tagged primers and high-temperature RT are required to eliminate self-priming of ZIKV RNA. To further improve the utility of the strand-specific assay, we developed fluorescent probes which allowed for simultaneous detection of positive-and negative-strand viral RNAs, with an internal control. This multiplexed RT-qPCR assay is both more sensitive and specific than previously-published assays for *Flaviviridae* strand-specific RNA detection. Using this assay, we demonstrate that negative-strand ZIKV RNA can be detected in both mammalian and mosquito cells, as early as 3-6 h post-infection in cell culture.

## 2. Materials and methods

### 2.1 In vitro transcription – standard curve generation

An infectious cDNA of ZIKV strain PRVAC59 (ZIKV^PR^; Genbank accession: KX377337) was kindly provided by Young-Min Lee (Utah State University) (32). To generate the template for positive-strand RNA transcription, the ZIKV infectious cDNA was linearized with BarI (Sibenzyme), verified by agarose gel electrophoresis, and column-purified using the Zymo DNA clean & concentrator kit (Zymogen) following the manufacturer’s instructions. To generate the negative-strand RNA IVT template, a T7 promoter was added to the negative-strand of the ZIKV infectious cDNA by PCR with Q5 DNA polymerase (New England Biolabs (NEB)) using primers T7-5’ZIKV(-)strand-FOR (5’-TAA TAC GAC TCA CTA TAG AGA CCC ATG GAT TTC CCC-3’) and 3’ZIKV(-)strand-REV (5’-AGT TGT TGA TCT GTG TGA ATC AG-3’). The PCR product was verified by agarose gel electrophoresis and PCR purified (Qiagen) prior to use as template in the IVT reaction. Two-hundred and fifty nanograms of linearized plasmid (positive-strand) or 150 ng PCR product (negative-strand) was used as a template in a run-off *in vitro* transcription reaction with SP6 RNA polymerase (NEB; positive-strand) or T7 RNA polymerase (NEB; negative-strand) following the manufacturer’s instructions with final NTP concentration of 1 mM. The *in vitro* transcribed RNA was treated with DNase I (NEB) and analyzed by agarose gel electrophoresis prior to purification with the Zymo RNA clean & concentrator kit (Zymogen) and the concentration was determination by UV-Vis spectrophotometry at 260 nm (Nanodrop).

### 3.2 Primer design

In order to facilitate strand-specific detection in a multiplex assay, we designed two sets of tagged primers with separate hydrolysis probes, which would enable us to detect both positive- and negative-strands simultaneously from the same sample, with an internal control. Tagged primers for the amplification of the negative-strand were chosen based on a previously published ZIKV RT-qPCR assay (26), and modified to match the nucleotide sequence of Asian lineage ZIKV isolates. Primers for the amplification of the positive-strand were selected such that their melting temperature was similar to those used for negative-strand detection, the amplicon was only present on genomic RNA (and not subgenomic flavivirus RNA), and that it was separated from the negative-strand by several kilobases (kb) so as to ensure specificity of probe binding to each RT product. The tag added to the positive-strand primers was adapted from (17). Mixed bases were included as needed to ensure primers were complementary to multiple ZIKV isolates. For the internal control (Glyceraldehyde 3-phosphate dehydrogenase (GAPDH)), primers were modified from (33) such that their melting temperature was similar to the ZIKV primers. Primers for mosquito GAPDH detection were designed with IDT’s PrimerQuest tool based on *Aedes aegypti* glyceraldehyde-3-phosphate dehydrogenase mRNA (XM_011494724.2; XM_019687453.2) All qPCR probes were designed using IDT’s PrimerQuest tool. All primer/probe sequences (listed in **Tables 1 and 2)** were checked for self-complementarity and potential heterodimerization using IDT’s oligoanalyzer tool.

**Table 1:**
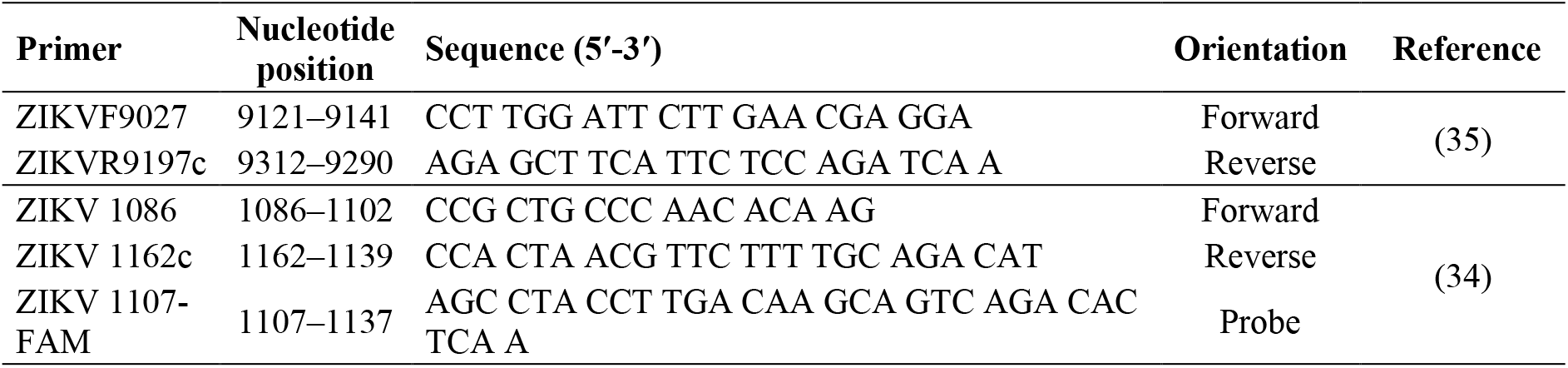
Primers used in standard RT-qPCR.

**Table 2:**
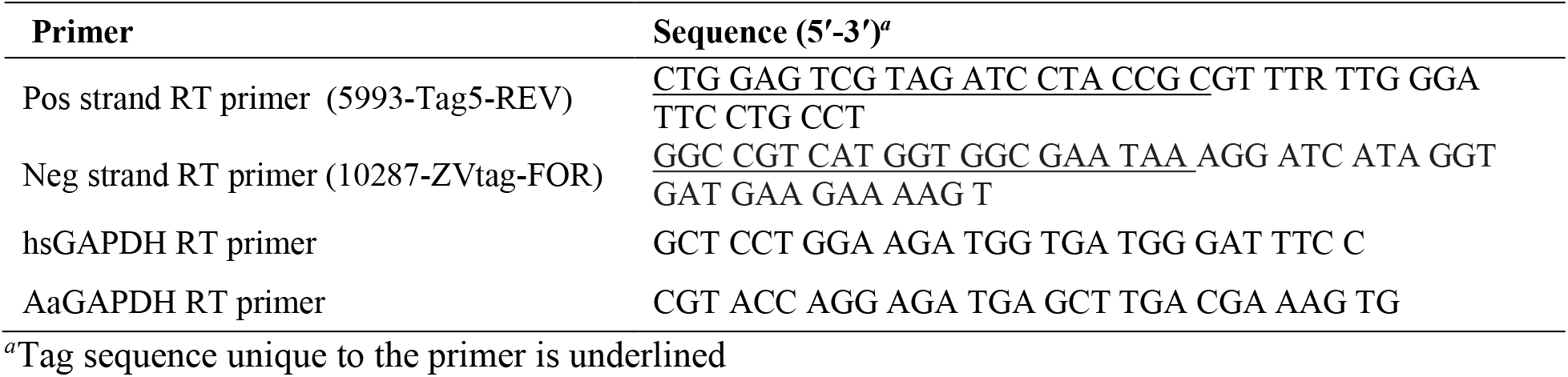
Tagged primers used in RT.

### 2.3 Standard RT-qPCR

RNA was mixed with 2 pmol of each primer (**Table 1, Figure 1**) and 0.5 mM dNTPs, denatured at 95 °C for 5 min and then immediately transferred to ice, where the RT buffer, DTT, RNase inhibitor (SuperasIN, Invitrogen or Ribolock, Thermo Scientific), and 0.5 µL SuperScript III reverse transcriptase (Invitrogen) were added as per the manufacturer’s instructions. RNA was then incubated at 55 °C for 30 min and the reaction was heat-inactivated at 70 °C for 15 min.

**Figure 1.**
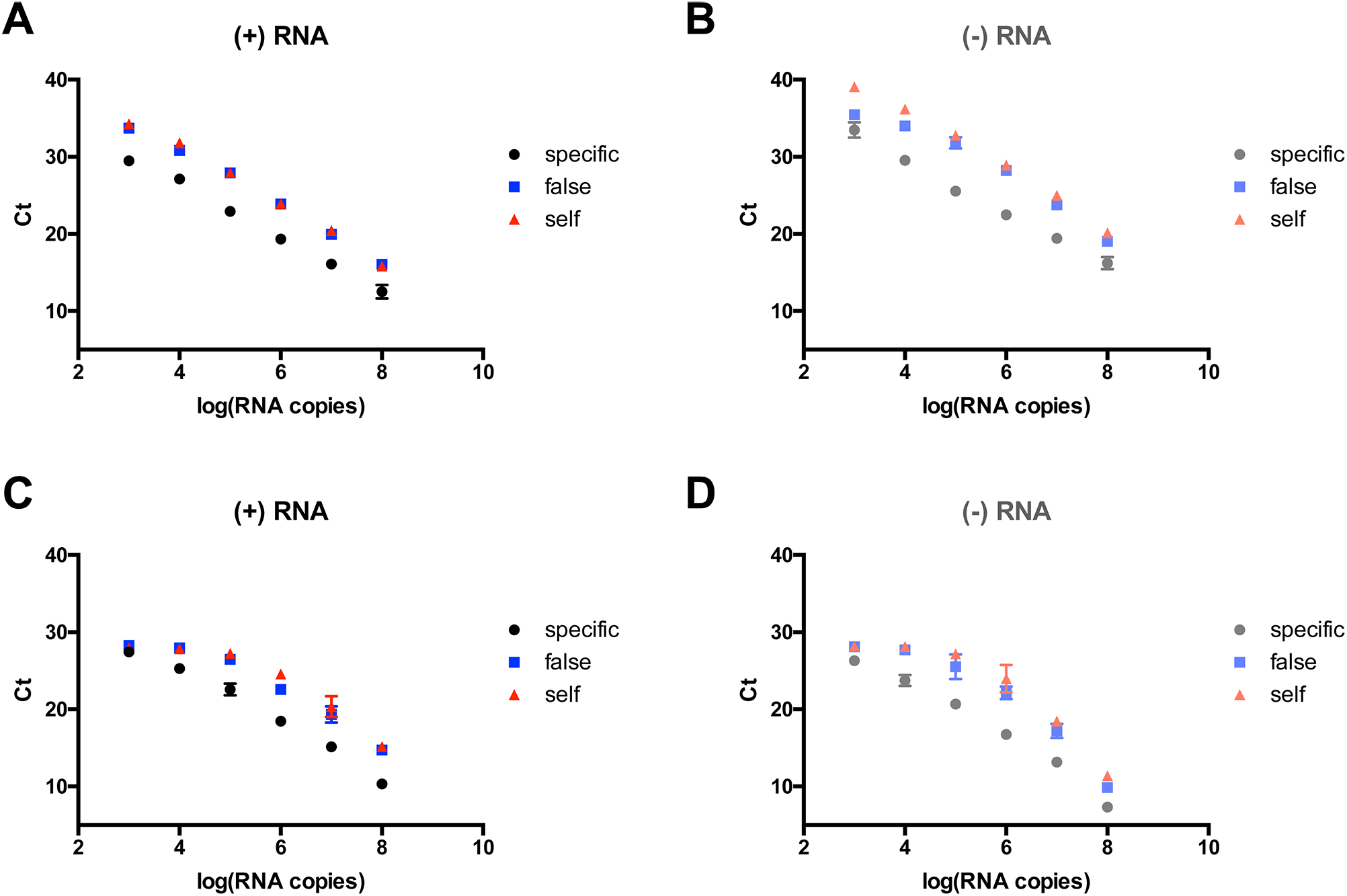
False- and self-priming is common with published ZIKV qPCR primer sets. Ten- fold serial dilutions of 10^8^ copies of positive-strand (A, C) or negative-strand (B, D) *in vitro* transcribed ZIKV RNA was analyzed by standard RT-qPCR using previously published ZIKV qPCR primer sets (Lanciotti *et al* 2008 (A, B) or Balm *et al* 2012 (C, D)). Specific, self-and false-priming were evaluated as described in the main text. The Ct value is plotted against the log of the RNA copy number (mean ± SEM, n = 2). Data points where amplification did not occur are not displayed on the graph.

Ten percent of the cDNA volume was used in the qPCR reaction. Quantitative PCR of the samples reverse transcribed using primers from Lanciotti *et al*. was performed using iTaq universal probes supermix (BioRad) with 300 nM each of the primers and probe (34). Thermocycling conditions were: 95ºC 3 min, followed by 40 cycles of: 95ºC 15 sec, 60ºC 60 sec + plate read. Quantitative PCR of the samples reverse transcribed with primers from Balm *et al*. was performed using SuperScript III Platinum SYBR Green One-Step qRT-PCR Kit (Invitrogen) with 200 nM primers and omitting the RT step (35). Thermocycling conditions were: 94 ºC 2 min, followed by 40 cycles of: 94 ºC 15 sec, 60 ºC 30 sec, 68 ºC 15 sec + plate read, followed by a Meltcurve from 65 ºC to 95 ºC. All qPCR reactions were performed on a CFX96 thermocycler (BioRad).

### 2.4 Strand-specific RT-qPCR

RNA was mixed with 10 nM each RT primer (**Table 2**) and 5 µM dNTPs, denatured at 95 °C for 5 min then immediately transferred to ice, where the RT buffer and RNase inhibitor (SuperasIN, Invitrogen or Ribolock, Thermo Scientific) were added. RNA was then transferred to 60 °C, after which 0.5 µL Maxima H-minus reverse transcriptase (Thermo Scientific) was added and the temperature was immediately transferred to 65 °C for 30 min. The RT reaction was heat-inactivated at 85 °C for 5 min and the cDNA was purified using the Zymo DNA clean & concentrator kit (Zymogen) according to the manufacturer’s instructions for cDNA clean-up. Quantitative PCR was performed using iTaq universal probes supermix (BioRad) with primer concentrations listed in **Table 3**.Thermocycling conditions were: 95 ºC 2 min, followed by 45 cycles of: 95 ºC 15 sec, 64 ºC 30 sec + plate read. For RNA extracted from mosquito cells, thermocycling conditions were: 95 ºC 2 min, followed by 45 cycles of: 95 ºC 15 sec, 64 ºC 60 sec + plate read to improve mosquito GAPDH PCR efficiency. ZIKV RNA was quantified based on a standard curve of *in vitro* transcribed positive-and negative-strand genomes, and was normalized to GAPDH by a modified ΔCt method (36).

**Table 3:**
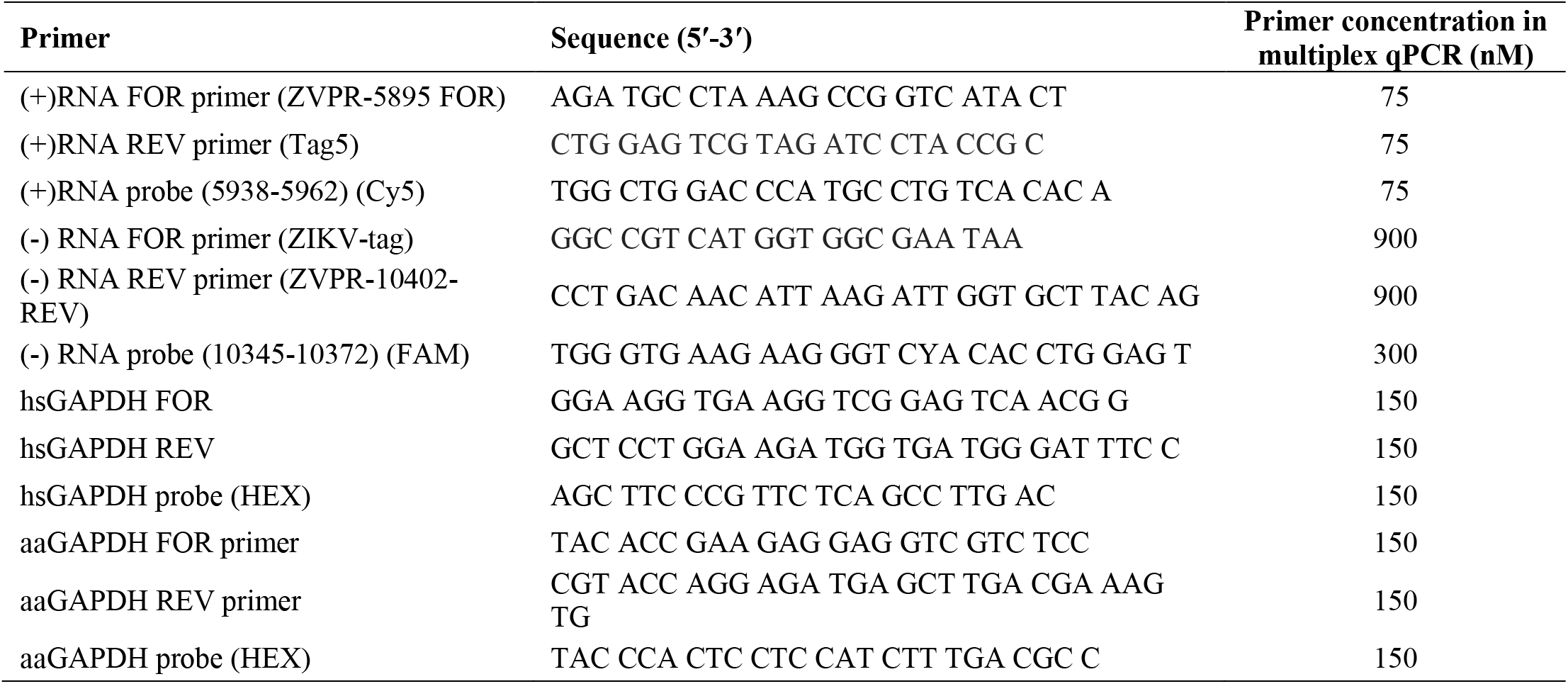
Primers used in multiplex strand-specific qPCR.

### 2.5 Cell culture

Human lung carcinoma (A549) cells, kindly provided by Russell Jones (Van Andel Institute, Michigan, U.S.A.), were maintained in Dulbecco’s modified Eagle’s medium (DMEM, Wisent Inc.) supplemented with 10% fetal bovine serum (FBS, Wisent Inc.), 1% Non-essential amino acids (Wisent Inc.), 1% L-glutamine (Wisent Inc.), and 1% penicillin/streptomycin (Wisent Inc.) at 37 °C/5 % CO_2_. Human choriocarcinoma (JEG-3) cells, kindly provided by Eric Miska (University of Cambridge, Cambridge, U.K.), were maintained in Eagle’s minimum essential medium (EMEM; Wisent Inc.) supplemented with 10% FBS, 1% Non-essential amino acids, 1% L-glutamine, 1% penicillin/streptomycin, and 1 mM sodium pyruvate (Wisent Inc.) at 37 °C/5% CO_2_. *Aedes albopictus* (C6/36) cells (ATCC) were maintained in Eagle’s minimum essential medium (EMEM; Wisent Inc.) supplemented with 10% FBS, 1% Non-essential Amino acids, 1% L-glutamine, 1% Pen/Strep, and 15 mM HEPES (Wisent Inc.) at 28 °C/5 % CO_2_.

### 2.6 Virus infections

An infectious cDNA of ZIKV strain PRVAC59 (Genbank accession: KX377337) was kindly provided by Young-Min Lee (Utah State University) (32). Viral stocks were generated by transfection of Vero cells with *in vitro* transcribed ZIKV RNA as previously described, followed by a single passage in Vero cells (37). Viral stocks were diluted to the indicated MOI in EMEM and were allowed to bind to subconfluent monolayers of cells for 1 h at 37 °C/5 % CO_2_, after which the inoculum was removed, cells were washed once with PBS, and media was replaced with fresh media containing 15 mM HEPES and 2% FBS. At the indicated time points post-infection, RNA was harvested in TriZol (Invitrogen) and extracted following the manufacturer’s instructions. Five-hundred nanograms of total RNA was used in the multiplex strand-specific RT-qPCR protocol.

## 3. Results

### 3.1 Conventional RT-qPCR is not strand-specific

We performed standard two-step RT-qPCR on *in vitro* transcribed ZIKV RNA positive-and negative-strand RNA using two previously published and widely used ZIKV primer sets (**Figure 1** and **Table 1**) (34, 35). We used different primers in the reverse transcription reaction to evaluate the potential contributions of false- and self-priming on the specific signal generated with the correct RT primer to detect the indicated strand. For specific priming, the reverse primer was used to reverse transcribe the positive-strand viral RNA, and the forward primer was used to reverse transcribe the negative-strand viral RNA. For false priming, the forward primer was used to reverse transcribe the positive-strand viral RNA, and the reverse primer was used to reverse transcribe the negative-strand viral RNA. For self-priming, no primers were added to the reverse transcription reaction. For both positive- and negative-strand viral RNA, both self- and false-priming occurred and was detectable when even as few as 10^3^ RNA copies were added to the RT reaction (**Figure 1**). Notably, there did not appear to be a difference in the degree of incorrectly primed cDNA products between the two chosen primer sets, even though they anneal to different regions of the viral genome. These results indicate that conventional RT-qPCR using several published ZIKV primer sets yields incorrectly-primed cDNA products and is therefore not suitable for quantitative strand-specific detection of viral RNAs.

### 3.2 Tagged primers with modified RT conditions largely eliminate false- and self-priming

We next tested whether the use of tagged primers would improve specificity. We designed tagged primers to specifically detect both positive- and negative-strand ZIKV RNA (**Figure 2A** and **Tables 2 and 3**, see **Materials and methods**). Although tagged primers have been widely reported to improve strand specificity, we found that tagged primers used alone in a standard RT-qPCR set-up did not sufficiently eliminate self- and false-priming of cDNA products (**Figure 2B-C**). To improve specificity, we purified cDNA products prior to qPCR analysis to remove any excess primer. We also increased the RT temperature, which required the use of a thermostable reverse transcriptase. We found that these conditions completely eliminated self-priming and greatly reduced the occurrence of false-priming (**Figure 2D-E**).

**Figure 2.**
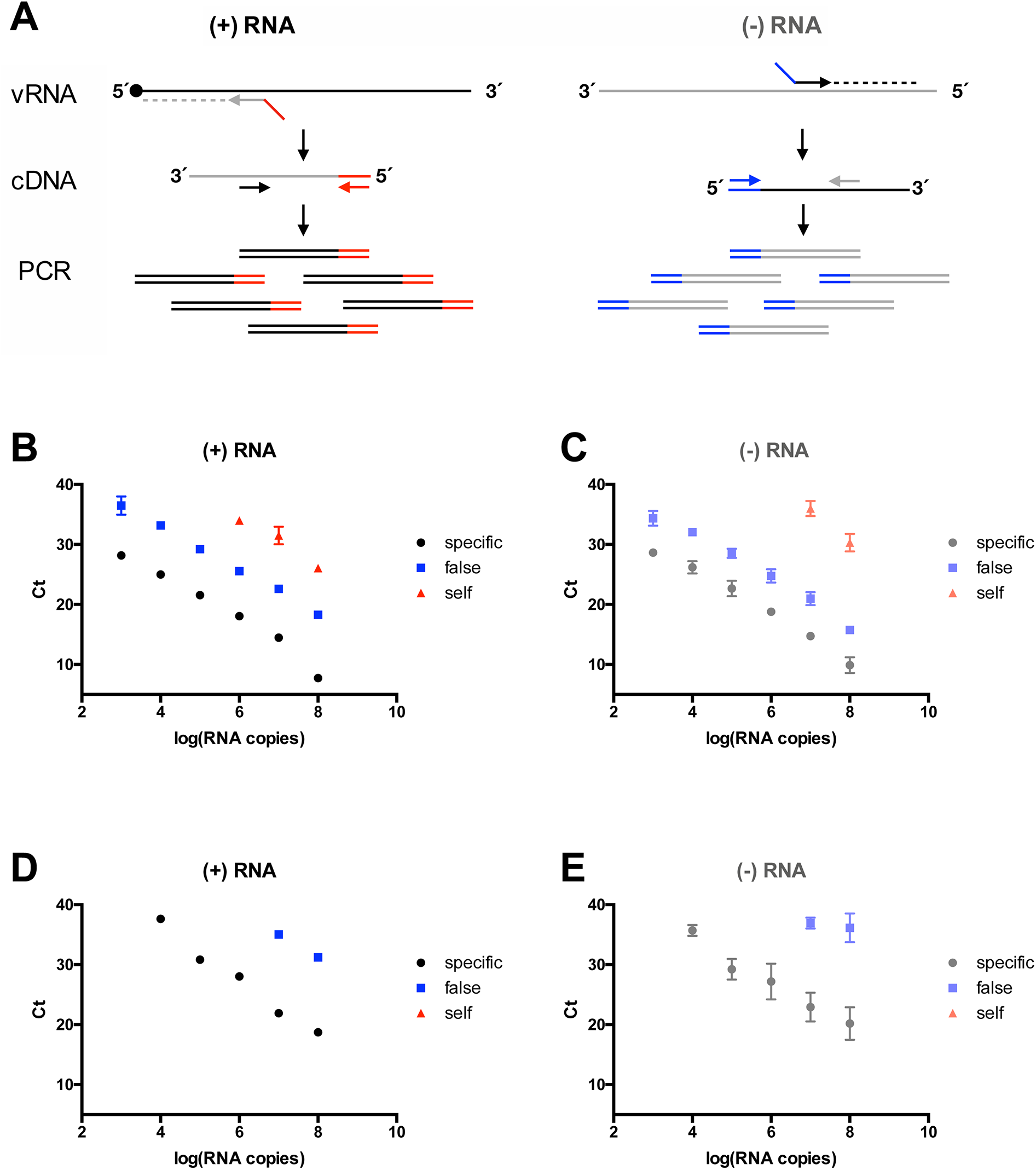
Tagged primers with modified RT conditions largely eliminate false- and self-priming. (A) Tagged primer strategy for strand-specific detection of positive- and negative-strand viral RNA (vRNA). Ten-fold serial dilutions of 10^8^ copies of positive-strand (B) or negative-strand (C) *in vitro* transcribed ZIKV RNA was reverse transcribed with SuperScript III Reverse transcriptase and analyzed by qPCR using tagged primers. Alternately, 10^8^ copies of positive-strand (D) or negative-strand (E) *in vitro* transcribed ZIKV RNA was serially diluted 10-fold and reverse transcribed with Maxima H minus Reverse transcriptase using tagged primers. cDNA was purified prior to analysis by qPCR as described in *Materials and Methods*. Specific, false, and self-priming were analyzed as described in **Figure 1**.The Ct value is plotted against the log of the RNA copy number (mean ± SEM, n = 2). Data points where amplification did not occur are not displayed.

### 3.3 Multiplex qPCR optimization

We next wanted to develop qPCR primers and probes which would enable us to multiplex detection of positive- and negative-strand viral RNAs together with an internal control (see **Materials and methods**). Given that there is reported to be up to 100-fold excess of positive-strand compared with negative-strand viral RNA during ZIKV infection, we first determined whether the primer concentrations needed for accurate detection of the negative-strand in multiplex PCR would need optimization (3-6). In a multiplex qPCR assay, it is important to verify that the amplification of high-abundance targets does not interfere with detection of low-abundance targets by depleting reagents (e.g. dNTPs, polymerase) at early cycles. Indeed, we found that when strand-specific cDNA from ZIKV-infected cells was subject to multiplex qPCR using typical qPCR primer concentrations (300 nM) for both positive- and negative-strand detection, detection of negative-strands was severely impaired (**Figure 3A**). This problem can be circumvented by modifying the primer concentration of high-abundance targets such that primers become limiting for that target (**Figure 3A**). The optimal primer concentrations necessary for accurate multiplex qPCR detection of all targets is provided in **Table 3**.

**Figure 3.**
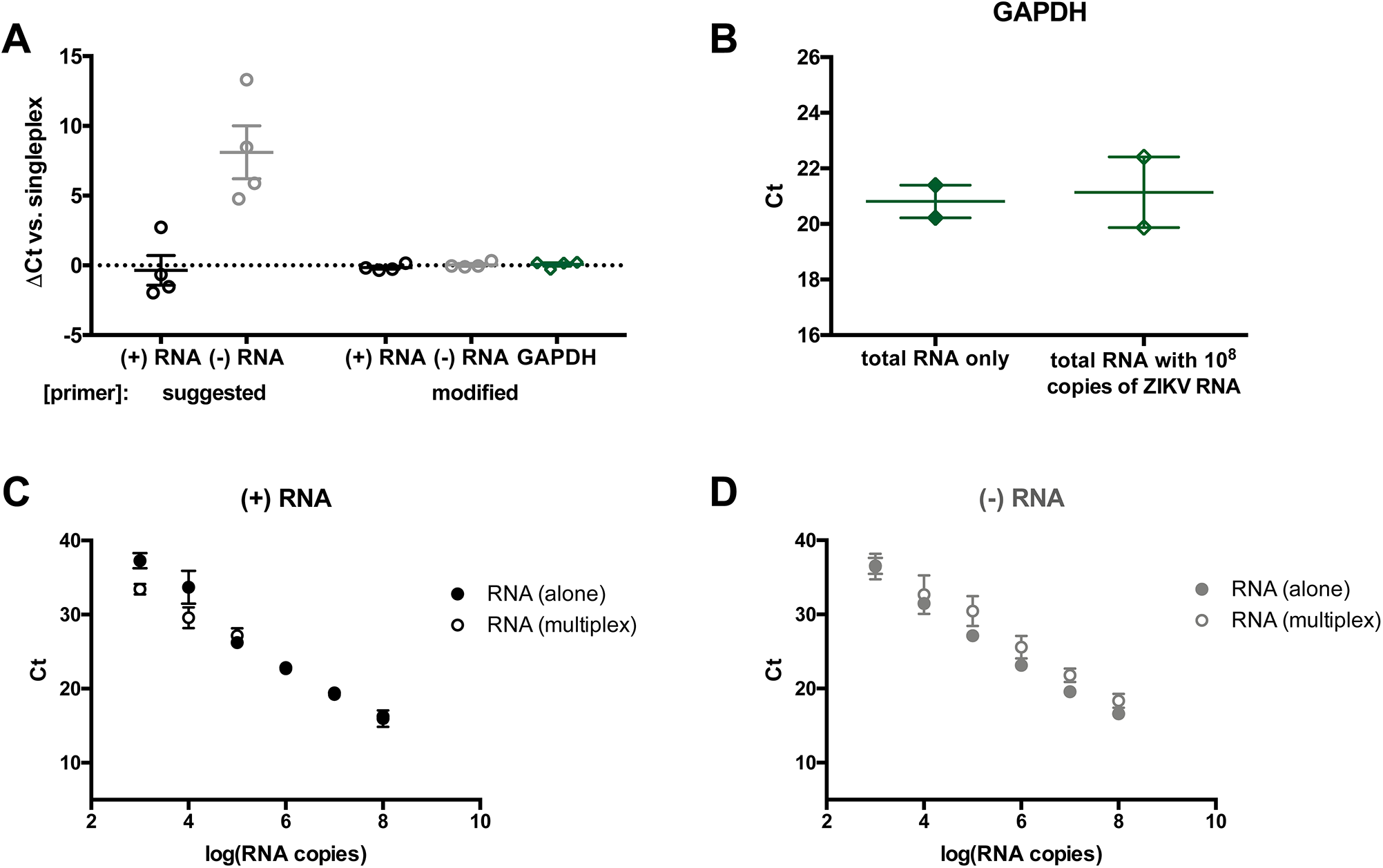
Multiplex qPCR optimization. (A) Total RNA from A549 cells infected with ZIKV was reverse-transcribed with positive- and negative-strand ZIKV tagged primers and GAPDH reverse primer. cDNA was purified prior to analysis by singleplex qPCR for the indicated target, or by multiplex qPCR analysis using standard PCR primer concentrations (300 nM) or primer concentrations modified for multiplex qPCR as described in **Table 3**.The ΔCt of the multiplex qPCR assay relative to the singleplex qPCR reaction is shown (mean ± SEM, n = 4). (B) Two-hundred nanograms total RNA from A549 cells with or without 10^8^ copies of both positive- and negative-strand *in vitro* transcribed ZIKV RNA added was reverse-transcribed with Maxima H minus reverse transcriptase and GAPDH reverse primer (total RNA only) or with GAPDH reverse primer and positive- and negative-strand ZIKV tagged primers. cDNA was purified prior to analysis by qPCR analysis with GAPDH primers (total RNA only) or by multiplex qPCR analysis with the modified primer concentrations from panel (A) (mean ± SEM, n = 2). (C, D) Ten-fold serial dilutions of 10^8^ copies of the indicated *in vitro* transcribed ZIKV RNA was reverse-transcribed with Maxima H minus reverse transcriptase and the corresponding tagged primers (RNA alone). Alternately, ten-fold serial dilutions of 10^8^ copies of both positive- and negative-strand *in vitro* transcribed ZIKV RNA was mixed with 200 ng total RNA, reverse-transcribed with Maxima H minus reverse transcriptase and positive- and negative-strand ZIKV tagged primers and GAPDH reverse primer (RNA in multiplex). cDNA was purified prior to analysis by qPCR analysis with the corresponding primer pair (RNA alone), or by multiplex qPCR analysis with the modified primer concentrations from panel (A) (mean ± SEM, n = 2).

Moreover, we showed that the addition of ZIKV RNA does not affect amplification of the housekeeping gene used as internal control (GAPDH) (**Figure 3B**), and that the addition of total RNA does not affect ZIKV amplification (**Figure 3C-D**). Importantly, GAPDH PCR efficiency is similar to ZIKV PCR efficiency (see section 3.5), so a modified ΔCt method could be used for normalization to the internal control (36). Similar optimization was performed for detection of ZIKV RNAs and an internal control RNA in mosquito cells (**Figure S1**). Finally, we validated the choice of housekeeping gene by determining that ZIKV infection does not affect GAPDH expression in both human and mosquito cells (**Figure S2**). Overall, these results suggest that the multiplex RT-qPCR assay can be used to accurately quantify ZIKV RNA from infected cells.

### 3.4 Demonstration of strand-specificity of the multiplex RT-qPCR assay

To determine the strand-specificity of the multiplex RT-qPCR assay, we mixed *in vitro* transcribed ZIKV positive- or negative-strand RNA with 100-fold, 1,000-fold, or 10,000-fold excess of the opposite strand (**Figure 4**). We found that positive-strand ZIKV RNA detection was specific up to at least 100-fold excess negative-strand RNA, after which excess negative-strands interfered with positive-strand detection (**Figure 4A**). Negative-strand RNA detection was specific up to at least 1000-fold excess positive-strand RNA, beyond which excess positive-strands could be non-specifically detected by the negative-strand assay (**Figure 4B**).

**Figure 4.**
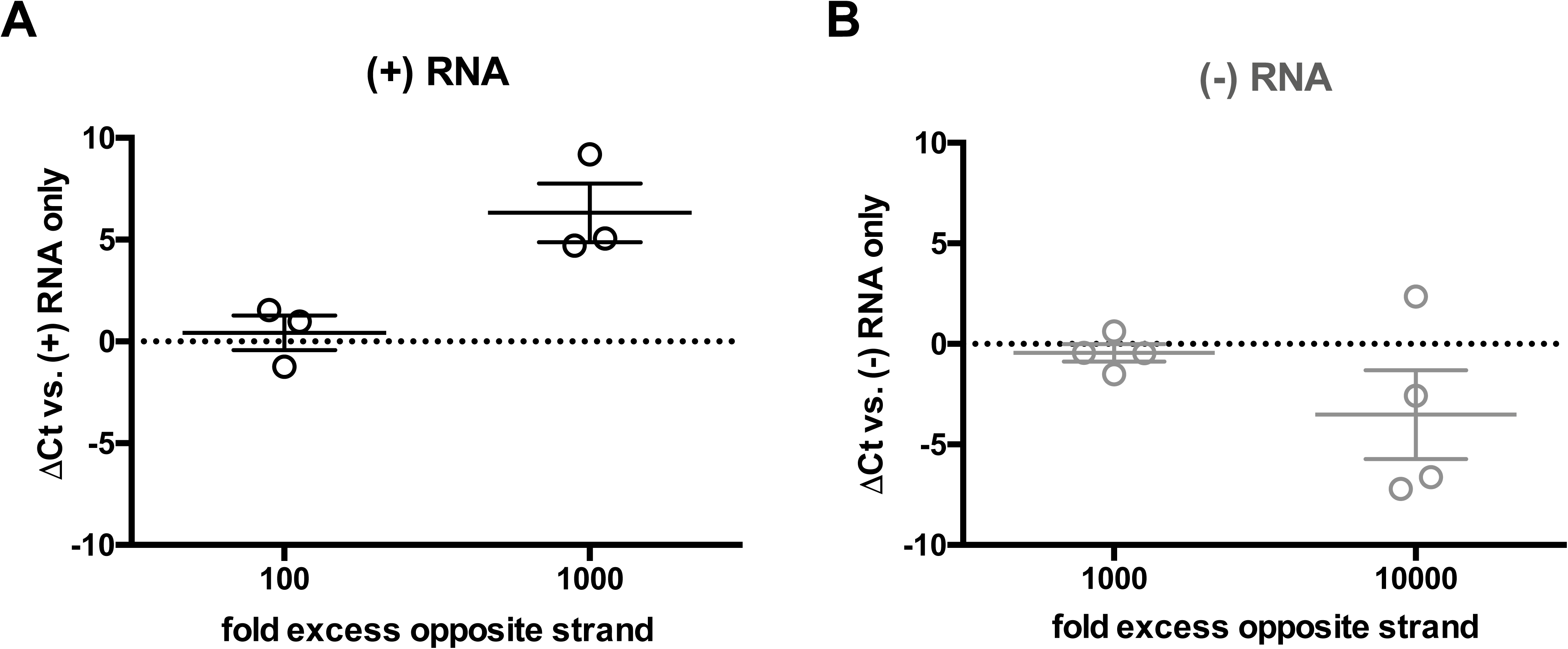
Specificity of strand-specific RT-qPCR assay. (A) 10^6^ or 10^5^ copies of *in vitro* transcribed positive-strand RNA mixed with 10^8^ copies of *in vitro* transcribed negative-strand RNA was analyzed by the strand-specific assay. The ΔCt relative to 10^6^ or 10^5^ copies of positive-strand RNA alone is shown (mean ± SEM, n = 3). (B) 10^5^ or 10^4^ copies of *in vitro* transcribed negative-strand RNA mixed 10^8^ copies of *in vitro* transcribed positive-strand RNA was analyzed by the strand-specific assay. The ΔCt relative to 10^5^ or 10^4^ copies of negative-strand RNA alone is shown (mean ± SEM, n = 4).

### 3.5 Validation of the strand-specific multiplex RT-qPCR assay

We next evaluated the sensitivity and reproducibility of the assay using *in vitro* transcribed positive-and negative-strand ZIKV RNA. A representative standard curve, consisting of 10-fold serial dilutions of 2.5 × 10^8^ copies each of positive-and negative-strands, was repeatedly subjected to the strand-specific multiplex RT-qPCR assay (**Figure 5**). Detection of positive-and negative-strands was similar across all dilutions. The standard deviation of each dilution was on average 1.73 Ct values for the positive-strand and 1.82 Ct values for the negative-strand, and did not vary across dilutions (see coefficient of variation (%CV) values in **Table 4**). The PCR efficiency averaged 92.9 ± 3.4% (R^2^ value of 0.980-0.999) and 90.1% ± 8.0% (R^2^ value of 0.989-0.998) for positive- and negative-strand detection, respectively. GAPDH PCR efficiency was 91.2 ± 5.8% (R^2^ value of 0.944-0.987). The lower limit of quantitation (LLOQ) ranged from 134 – 333 and 130 – 2109 copies per reaction for the positive- and negative-strand, respectively. Given that up to 500 ng RNA was analyzed per reaction, this represents a lower limit of quantitation of 0.27-0.67 and 0.26-4.22 positive- and negative-strand copies per ng RNA, respectively.

**Table 4:** Intra-assay variability.

| log (RNA copies) | 8 | 7 | 6 | 5 | 4 | 3 | 2 |
| --- | --- | --- | --- | --- | --- | --- | --- |
| (+) RNA (%CV)<sup>a</sup> | 6.7% | 10.4% | 10.0% | 7.0% | 6.3% | 5.5% | 4.3% |
| (-) RNA (%CV) | 4.9% | 10.1% | 9.1% | 6.3% | 7.1% | 6.3% | 5.2% |

<sup>a</sup>%CV = coefficient of variation calculated from Ct values (n = 5).

**Figure 5.**
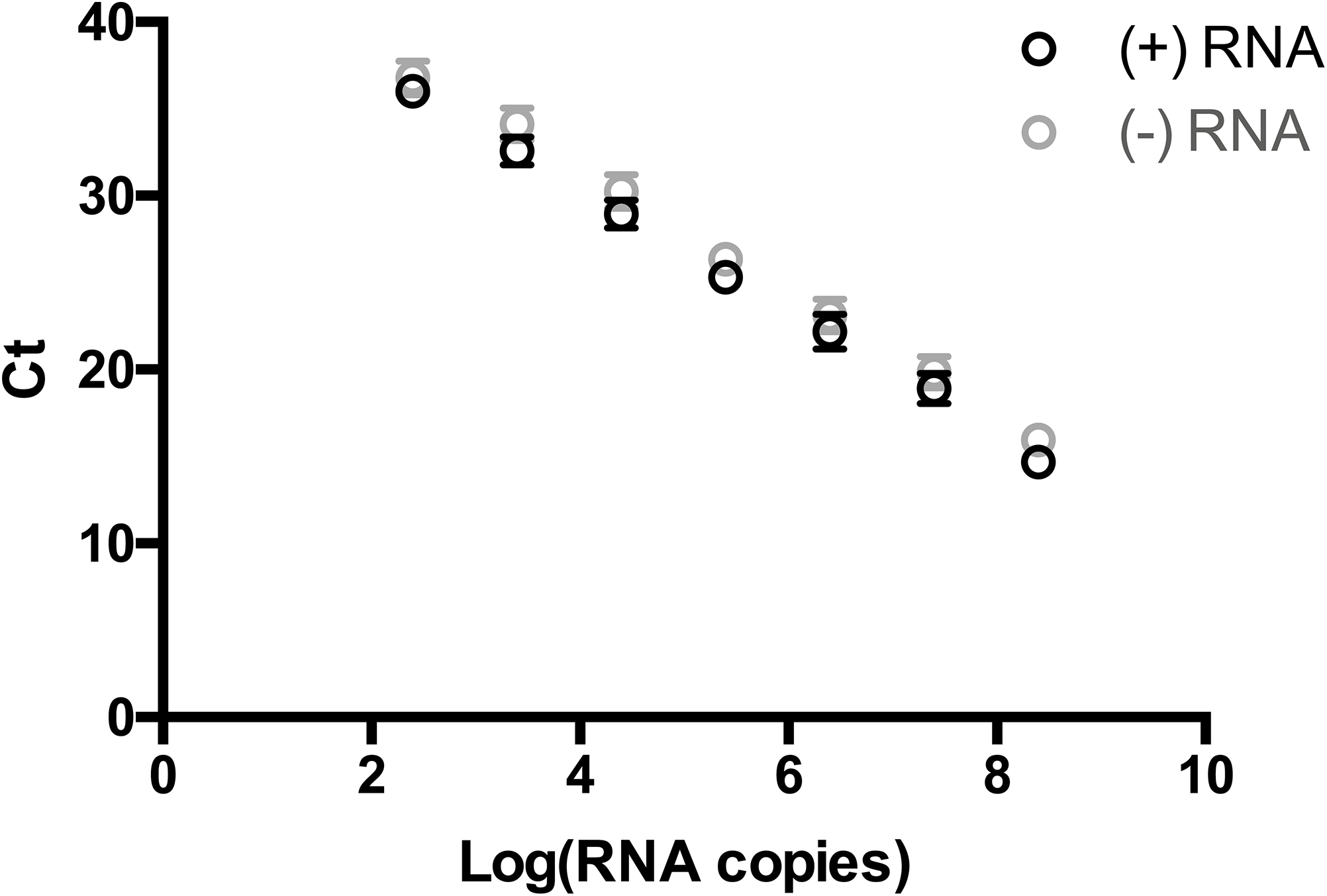
Reproducibility of strand-specific RT-qPCR assay. Ten-fold serial dilutions of 2.5 × 10^8^ copies each of *in vitro* transcribed ZIKV positive-strand and negative-strand RNA was analyzed by the strand-specific RT-qPCR assay. Average Ct value is plotted against the log of the RNA copy number (mean ± SEM, n = 5).

### 3.6 Quantitation of positive- and negative-strand ZIKV RNA in cell culture

In order to understand ZIKV replication dynamics in cell culture, the strand-specific RT-qPCR assay was used to quantify ZIKV RNA in infected cells. We performed one-step kinetics in human placental choriocarcinoma (JEG-3) cells infected with ZIKV (**Figure 6A**). Interestingly, we found that negative-strand RNA could be detected as early as 3 h post-infection. Correspondingly, positive-strand RNA began to increase between 3 and 6 h post-infection, again implying that negative-strands were synthesized by this time point. Both positive- and negative-strand RNA continued to increase throughout the duration of experiment, indicating that active viral replication was occurring. The (+):(-) RNA ratio was approximately 30-60:1 at all timepoints where negative-strand RNA could be quantified.

**Figure 6.**
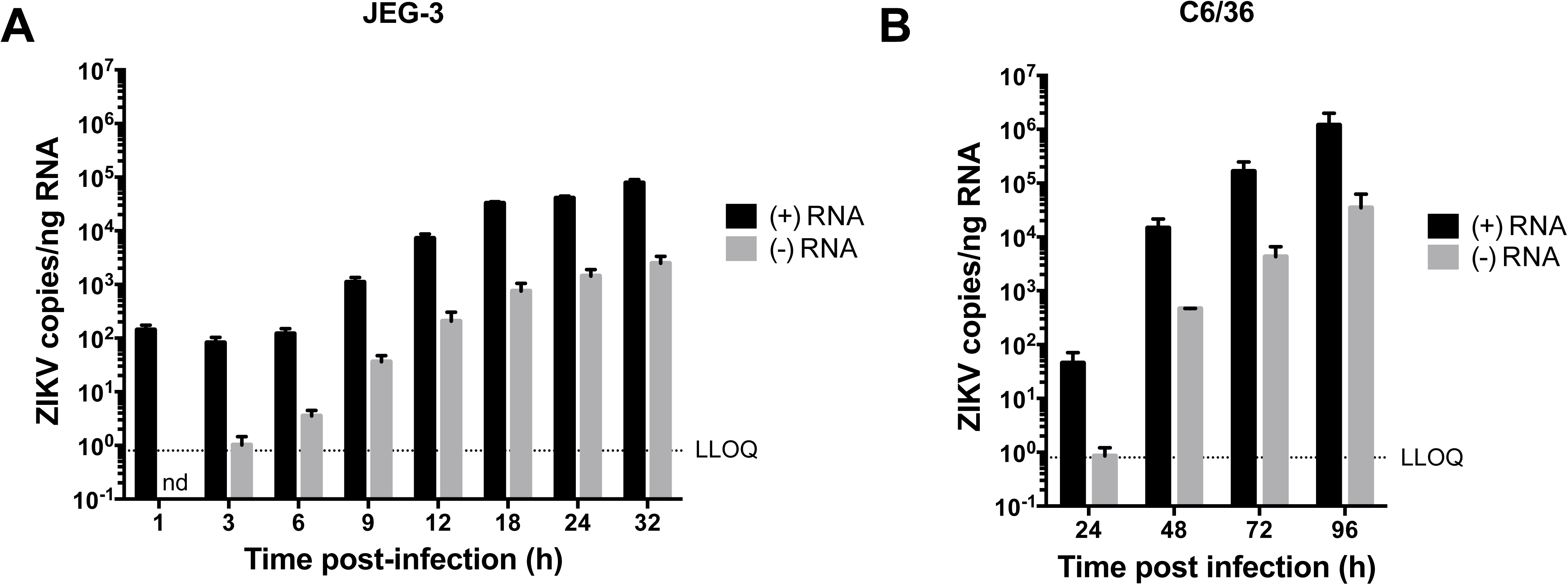
Representative results in mammalian and mosquito cells. (A) JEG-3 cells were infected with ZIKV^PR^ (MOI 3) and RNA was harvested at the indicated time points. Positive- and negative-strand viral genomes were quantified by the strand-specific RT-qPCR assay and normalized to GAPDH (mean ± SEM, n = 3). (B) C6/36 cells were infected with ZIKV^PR^ (MOI 0.01) and RNA was harvested at the indicated time points. Positive- and negative-strand viral genomes were quantified by the strand-specific assay, normalized to GAPDH (mean ± SEM, n = 2). nd; not detected.

Similarly, active viral replication was also detectable in C6/36 mosquito cells (**Figure 6B**). We found that ZIKV positive- and negative-strand RNAs increased throughout the duration of the experiment, suggesting that active viral replication was occurring. Interestingly, even at very low MOIs (i.e. MOI = 0.01), negative-strand RNA could be detected as early as 24 h post-infection. In mosquito cells, the (+):(-) RNA ratio was approximately 30-50:1. Overall, our results demonstrate that the strand-specific RT-qPCR assay can be used to quantify ZIKV positive- and negative-strand RNA from both human and mosquito cells.

## 4. Discussion

Herein, we show that incorrectly-primed cDNA products are generated during reverse transcription with commonly-used ZIKV primer sets. Consequently, we developed a strand-specific multiplexed RT-qPCR assay for the quantitation of ZIKV RNA in infected human and mosquito cells. We show that the assay provides sufficient specificity for detection of positive-and negative-strand ZIKV RNA during infection. Finally, we demonstrate that the assay can be multiplexed and used to quantify ZIKV RNA replication in both human and mosquito cells. In summary, the strand-specific RT-qPCR assay developed herein is a useful tool for the evaluation of ZIKV RNA replication kinetics, tropism, and persistence in both human and mosquito cells.

Interestingly, the majority of the nonspecific priming observed during conventional RT-qPCR was due to self-priming of the viral RNA. There is a high degree of secondary structure observed throughout the ZIKV genome, which may contribute to self-primed cDNA synthesis (38, 39). The degree of self-priming did not differ between the two primer sets, despite their specificity for different regions of the viral genome. This suggests that self-priming occurs throughout the ZIKV genome, and should be considered when performing RT on ZIKV RNA. Furthermore, the genomes of diverse RNA viruses are highly structured, and as such self-priming is likely to be a problem for strand-specific RNA detection of multiple viral families (40-42). Indeed, self-priming during RT has been demonstrated to occur for numerous RNA viruses, including Dengue virus, hepatitis E virus, human rhinovirus, hepatitis C virus, and several plant RNA viruses (16, 43-46).

Importantly, use of tagged primers with high-temperature RT completely eliminated self-priming, which constituted the majority of nonspecific events. Despite this, the limit of strand-specificity for negative-strand detection was 1000-fold excess positive-strands. This is likely due to small degree of false priming that could still be detected even with tagged primers. Interestingly, although false priming of the positive-strand occurred to a similar degree, 1000-fold excess negative-strand decreased rather than increased signal, suggesting that the excess negative-strand impeded positive-strand detection, likely through hybridization with complementary RNA. Other strand-specific RT-qPCR assays using tagged primers have reported complete specificity (i.e. no detection of even very high amounts of the incorrect strand), suggesting that further modification of our tagged primers has the potential to further improve specificity. However, higher specificity likely comes as a trade-off to sensitivity (7). Increased specificity, i.e. beyond 1000:1 (+):(-) RNA, is likely unnecessary as the (+):(-) RNA ratio during infection is within our assay’s range of specificity, where a ratio of 10:1-100:1 excess positive-strand RNA has been reported (3-6). Furthermore, decreased sensitivity would impede negative-strand detection at early time points and render the assay less able to quantify early events in viral replication.

Although both viral RNA targets were detected similarly, PCR efficiency was more variable in the negative-strand assay than the positive-strand assay. This was likely due to stochastic detection of the lowest concentration of negative-strand RNA standards (47), as indicated by the greater degree of variability of the negative-vs. positive-strand LLOQ, rather than overall greater variability of negative-strand detection as the standard deviation of each Ct did not vary between (+) and (-) RNA. PCR efficiency between 90-110% is generally considered acceptable, and both positive- and negative-strand detection fell within this range (48). It is possible that this variability in efficiency of negative-strand detection was caused by the quality of the *in vitro* transcribed RNA transcripts used in the experimental determination of PCR efficiency.

Despite the above-described limitations, to our knowledge this assay represents the first validation of a strand-specific multiplex RT-qPCR assay for ZIKV RNA quantitation. The assay presented herein improves over similar previously validated *flavivirus* strand-specific RT-qPCR assays, which are not sensitive enough to quantify early events in viral replication (7). Depending on the amount of input RNA added to the RT reaction, the lower limit of quantitation of ∼200-2000 copies/reaction can be as low as 0.5-5 copies per ng input RNA. Estimates suggest that a single mammalian cell contains approximately 10-20 pg total RNA (49); as such, the lower limit of detection of our assay is less than one copy of viral RNA per cell. Thus, this enables the study of very early events in the viral life cycle.

Mosquitoes are an important component of the ZIKV transmission cycle and therefore the study of viral replication in mosquitoes may provide insights into potential strategies to block transmission. To our knowledge, detection of ZIKV negative-strand RNA from either mosquito cells in culture or live mosquitoes is rare (50). Nonetheless, the assay developed herein allows quantification of ZIKV replication in mosquito cells and could potentially be used to quantify viral replication from mosquito tissues, which would expand our knowledge of transmission bottlenecks and mechanisms of viral replication in the insect vector.

Overall, this study adds to the growing body of evidence which suggests that conventional RT-qPCR methods do not accurately detect viral genomes in the presence of other forms of viral RNA (7, 8, 10, 51-53). Results from studies that present strand-specific RT-qPCR data in the absence of a validated strand-specific assay should therefore be interpreted with the caveat that one cannot necessarily conclude that RNA copy number represents uniquely genomic or antigenomic RNA. For viruses which make multiple mRNA species in addition to genomic and antigenomic RNA (e.g. Coronaviruses), conventional RT-qPCR is perhaps even less likely to provide accurate quantitation of viral genomic RNAs. Importantly, similar limitations of strand-specific detection are not limited to positive-sense RNA viruses (9, 51, 52). Nevertheless, with some thought it is possible to design strategies to specifically detect the RNA species of interest even in the presence of several types of viral RNA (51, 52, 54, 55).

## Acknowledgements

We thank Young-Min Lee (Utah State University) for providing ZIKV^PR^ infectious cDNA. We also thank Russell Jones (Van Andel Institute, Michigan, U.S.A.), and Eric Miska (University of Cambridge, Cambridge, U.K.) for providing human lung carcinoma (A549) and human choriocarcinoma (JEG-3) cells, respectively. Finally, we thank Quinn Abram for providing RNA from infected C6/36 cells.

## Author Contributions

T.R.B and S.M.S designed the study; T.R.B. and A.B.W performed the experiments and analyzed the data, and T.R.B. wrote and edited the manuscript with assistance from S.M.S. and A.B.W.

## Graphical Abstract

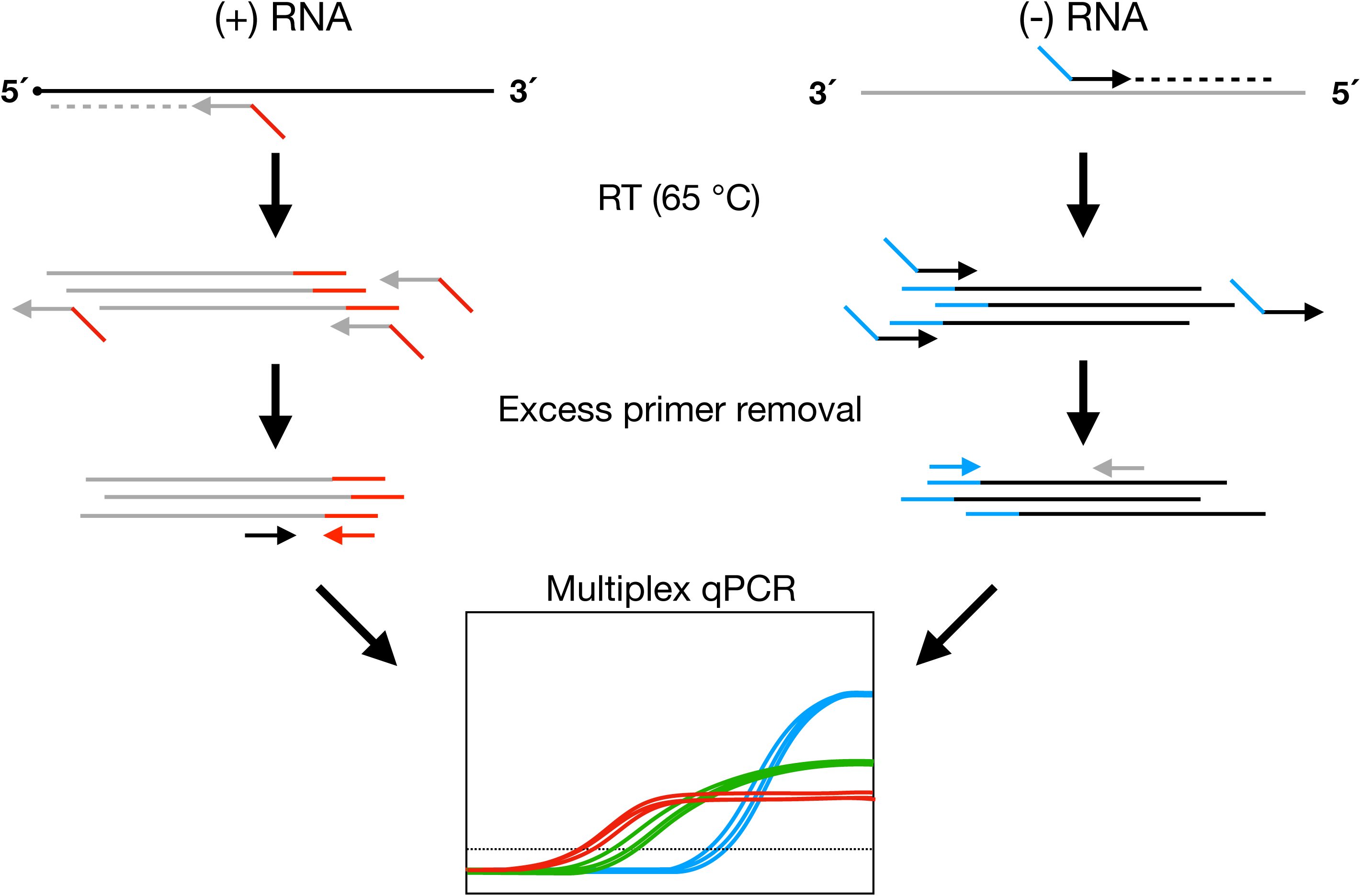

