## Supplemental Figures for "A highly sensitive strand-specific multiplex RT-qPCR assay for quantitation of Zika virus replication"

**SUPPLEMENTARY FIGURES**

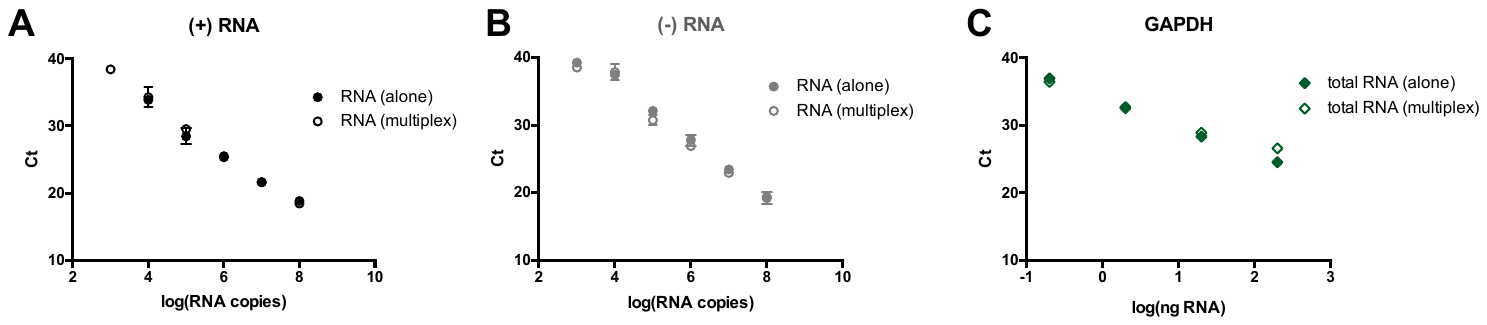

**Figure S1.** **Assay validation for mosquito cells.** (A, B) Ten-fold serial dilutions of 10^8^ copies of the indicated *in vitro* transcribed ZIKV RNA was reverse-transcribed with Maxima H minus reverse transcriptase and the corresponding tagged primers (RNA alone). Alternately, ten-fold serial dilutions of 10^8^ copies of both positive- and negative-strand *in vitro* transcribed ZIKV RNA was mixed with 200 ng total RNA from mosquito cells, reverse-transcribed with Maxima H minus reverse transcriptase and positive- and negative-strand ZIKV tagged primers and mosquito GAPDH reverse primer (RNA in multiplex). cDNA was purified prior to analysis by qPCR analysis with the corresponding primer pair (RNA alone), or by multiplex qPCR analysis with primer concentrations listed in **Table 3** (mean ± SEM, n = 2). (C) Ten-fold serial dilutions of 200 ng total RNA from mosquito cells was reverse-transcribed with Maxima H minus reverse transcriptase and mosquito GAPDH reverse primer (total RNA only) or with mosquito GAPDH reverse primer and positive- and negative-strand ZIKV tagged primers. cDNA was purified prior to analysis by qPCR analysis with GAPDH primers (total RNA only) or by multiplex qPCR analysis with the primer concentrations listed in **Table 3** (mean ± SEM, n = 2).

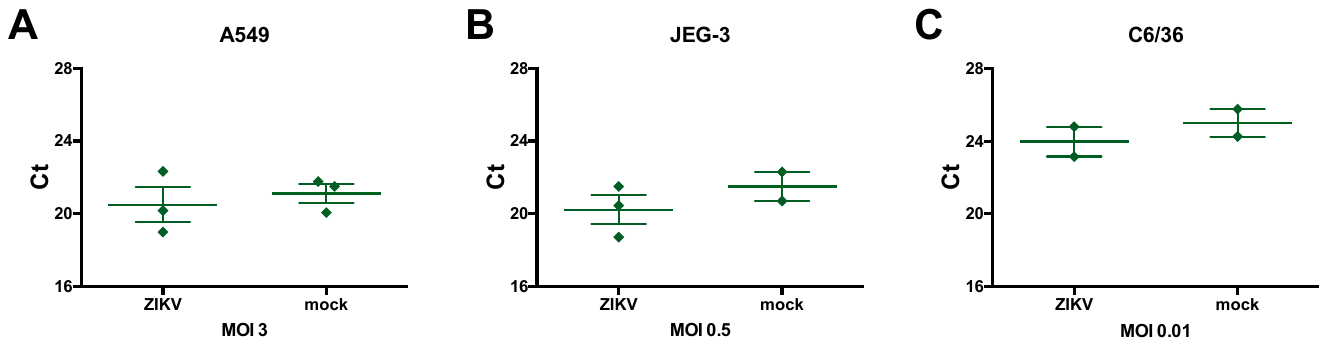

**Figure S2.** **ZIKV infection does not affect GAPDH expression.** Total RNA from mock-infected cells or cells infected with ZIKV was analyzed by the strand-specific RT-qPCR assay, and the resulting GAPDH Ct is shown. (A) A549 cells (MOI 3) at 24 h post-infection, (B) JEG-3 cells (MOI 0.5) at 24 h post-infection, or (C) C6/36 cells (MOI 0.01) at 72 h post-infection (mean ± SEM, n = 2-3).
